# Gene set inference from single-cell sequencing data using a hybrid of matrix factorization and variational autoencoders

**DOI:** 10.1101/740415

**Authors:** Soeren Lukassen, Foo Wei Ten, Roland Eils, Christian Conrad

**Affiliations:** Charité – Universitätsmedizin Berlin, Digital Health Center, Berlin, Germany; Berlin Institute of Health (BIH), Berlin, Germany; Health Data Science Unit, University Hospital Heidelberg, Heidelberg, Germany

## Abstract

Recent advances in single-cell RNA sequencing (scRNA-Seq) have driven the simultaneous measurement of the expression of 1,000s of genes in 1,000s of single cells. These growing data sets allow us to model gene sets in biological networks at an unprecedented level of detail, in spite of heterogenous cell populations. Here, we propose an unsupervised deep neural network model that is a hybrid of matrix factorization and conditional variational autoencoders (CVA), which utilizes weights as matrix factorizations to obtain gene sets, while class-specific inputs to the latent variable space facilitate a plausible identification of cell types. This artificial neural network model seamlessly integrates functional gene set inference, experimental batch effect correction, and static gene identification, which we conceptually prove here for three single-cell RNA-Seq datasets and suggest for future single-cell-gene analytics.

## Main

Gene expression is a hierarchical, structured and highly controlled process that is the major determinant of identity and state in cells. Thereby, genes of same functional ‘origin’ can typically be grouped into *sets* that share a common expression, regulation, or function across different cells. These *gene sets* or genetic pathways can be identified through prior knowledge. Additionally, co-expression and co-regulation of genes from measured experiments have been used to inform these pathways by regulators^1^ or tissue-specificty^2^. However, manual generation and curation of these data is biased towards published sets^3^, error-prone, and time consuming as well. Until recently, the modeling of gene sets using advanced machine learning (ML) approaches was limited by unbalanced RNA sequencing studies measuring the expression of 10,000s known genes expressed in only few tissue samples, completely neglecting cellular heterogeneity within bulk samples. With the advent of single-cell RNA sequencing the simultaneous measurement of individual gene expression in 1,000s to 100,000s single cells is feasible. By massive probing of gene expression correlation in the variation of gene expression within tissues, subtle genetic relationships in heterogenous tissues can now be studied.

Because of typical multiple assignments of genes (N) to different sets (M), methods that parameterize weights to connect genes to sets, capture biological complicity more accurately than any hard-clustering (1-to-M). To this end, in very high-dimensional scRNA-Seq factorizations and rotations of the gene expression matrices such as principal component analysis (PCA) or its variant singular value decomposition (SVD)^4–6^, and in case of multiple samples related canonical correlation analysis (CCA), are applied^7,8^. Easier interpretability of matrix factorization in scRNA-Seq has been achieved through enforcing sparsity of weights in sparse decomposition of arrays (SDA)^9,10^. Non-negative matrix factorization (NMF) and derived methods specifically account for the inherent non-negativity of gene expression by decomposing the expression matrix into matrices with strictly positive values^11^ and are accordingly performed in single cell sequencing analysis^12–14^. Moreover, further decomposition steps can adequately add hierarchical organization of the component spaces in NMF^15,16^. Although these deep NMF algorithms are algorithmically similar to deep autoencoders (AEs)^16,17^, they do not satisfy more complex, nonlinear relationships^18^. Variational autoencoders (VAE) estimating the underlying probability distribution of the input data^19,20^ have been successfully used in scRNA-Seq, with a special focus on the latent variable space for dimension reduction and clustering^21,22^. It is worth noting that the two previously mentioned algorithms are not mutually exclusive and concepts may be combined for specific applications (NMF^12^ and AE^12,17^), becoming more practical if the latent variable space takes specific cell type classes into account across training of the network model.

### A hybrid VAE and NMF based network for gene set inference

We propose a VAE and NMF based neural network architecture for gene set inference and batch effect correction using scRNA-Seq data as input, named **conditional variational autoencoder (CVA)**. This architecture can assign dimensions or latent variables to certain predefined classes by setting the activation of all other latent variables to zero. The resultant network has a shared encoder and decoder network, but a class-specific, spherical latent variable space (Fig. 1a). This allows for separation of the contribution of overlapping effects, e.g. when the latent space separation is used to encode for cell type, batch, and other confounders. Individual decomposing layers in the neural network can access different, hierarchically organized structures of the data. In the case of cell types as latent variable determinants, hierarchical grouping of pathways and co-expressed gene sets are modeled in decoder layers of the CVA (Fig. 1b). In contrast to most standard neural network architectures, we omitted the bias terms from all decoder layers, making our decoder network more similar to an NMF. Keeping the bias term of the output layer as trainable parameter abstracts static components of gene expression, i.e. housekeeping genes (expressed in all cell types). In contrast to algorithms performing matrix factorizations, we implement activation functions for all layers to mimic complexity in gene expression and toggle switching of e.g. transcription factors. In the context of our VAE, gene expression and pathway activity correspond to layer activations, which are consequently forced to be zero or positive by the rectifier linear unit (ReLU) activation (Fig. 1c). The separation of activation and weights allows the isolation of the magnitude and direction of the influence of gene sets on genes, irrespective of their current activation state. The connection strength between a hidden decoder layer and the output layer representing individual genes can thus be obtained by multiplying the weight matrices of all subsequent layers. This results in a single matrix of genes assigned to introduced cell types (Fig. 1d).

**Figure 1.**
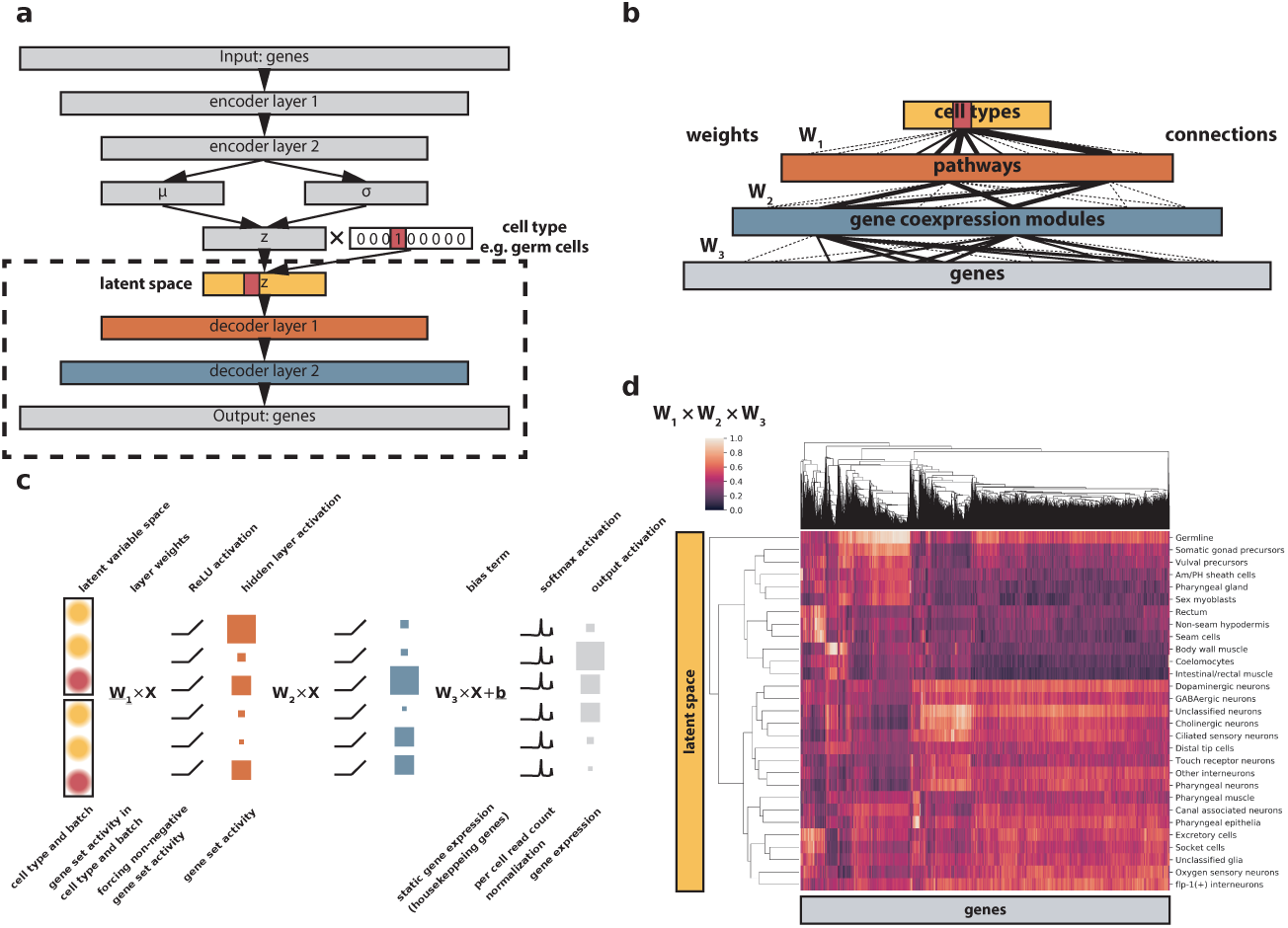
CVA network architecture. a) General architecture of the deep neural network employed here. b) Illustration of the connectivities and encoded entities within the decoder part of the network, which is used for gene set inference. c) Mathematical operations in the decoder network and their counterparts in biological systems. d) Clustered heatmap depicting the weight mappings from the class specific latent variable space (rows) to genes (columns).

### Gene set inference from entire organisms

In order to evaluate the performance of our CVA model for gene set inference, we applied the algorithm to a scRNA-Seq dataset from *C*. *elegans*, comprising over 30,000 cells of 27 different cell types^23^. This dataset captures most cell types present in the adult worm, making it ideally suited for gene set and pathway inference. In a first sanity check, we obtained the connection weights from the cell type encoding latent variable space to the output genes. We then calculated the associated genes and performed gene set enrichment analyses using Metascape^24^ as benchmarks. Example mappings for different cell types showed a good concordance of the obtained pathway enrichments with expected values (Fig. 2a-c). The presence of ‘acetylcholine metabolic process’ in the enriched terms for cholinergic nerve cells (Fig. 2a) shows that specific terms for subtypes are detected by the algorithm. We then set out to investigate gene sets activated by specific (mathematical) neurons of the hidden decoder layers (Fig. 2d-g). Early decoder layer neurons tended to access more complex pathways, such as molting (Fig. 2d), stress response (Fig. 2e), and muscle development (Fig. 2f), while neurons in late decoder layers access more specific gene sets, such as protein targeting to membranes (Fig. 2g). In conclusion, we successfully deciphered specific biological useful pathways and as well as co-expressed gene sets.

**Figure 2.**
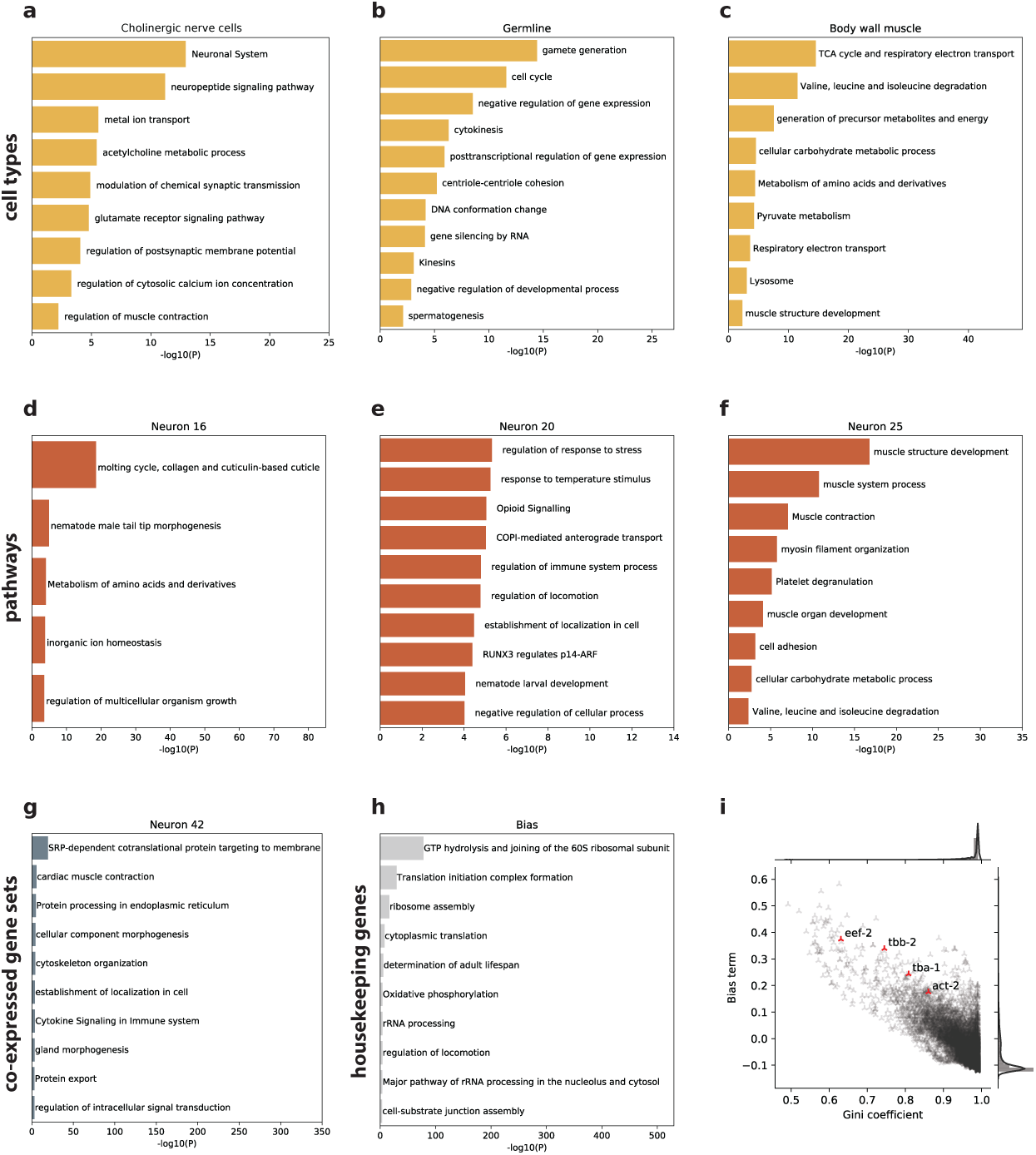
Gene set and housekeeping gene inference from *C*. *elegans* scRNA-Seq. a – c) Gene set enrichment of the mappings from three cell type specific latent variable spaces to genes. Only gene sets with an enrichment P-value < 10-2 are shown and lists are truncated to a maximum of 11 entries. a) cholinergic neuron, b) germ cell, c) body wall muscle specific latent variable space mapping. d – f) Gene set enrichment of weight mappings from three neurons from the first decoder layer to genes. g) Gene set enrichment of weight mappings from one neuron from the second decoder layer to genes. h) Gene set enrichment of the genes with high bias terms associated with them. i) Scatterplot and kernel density estimate of bias terms versus Gini coefficient in the normalized expression matrix for all genes in the dataset. The *C*. *elegans* orthologs of β-actin (act-2), α-tubulin (tba-1), β-tubulin (tbb-2), and the eukaryotic translation initiation factor 2 (eef-2) are highlighted. Colors of the graphics a)-h) correspond to the layers of the CVA in Fig. 1b.

### Housekeeping gene identification

To test whether the bias term of the output layer does indeed identify the static component of gene expression, which should correspond to housekeeping genes, we investigated the bias terms further. Pathway enrichment of genes that had exceedingly high bias terms associated with them revealed a strong link to housekeeping functions such as ribosome biogenesis and function, oxidative phosphorylation, and cytoskeletal biogenesis and function (Fig. 2h). Housekeeping genes should show very little variability in the input expression matrix and should thus have low Gini coefficients in the dataset used for training. To investigate the relationship between the bias terms and the Gini coefficients, we plotted both against each other and observed a strong negative correlation, as would be expected if bias terms capture the static element of gene expression (Fig. 2i). We highlighted the *C*. *elegans* orthologs of β-actin (act-2), α-tubulin (tba-1), β-tubulin (tbb-2), and eukaryotic translation elongation factor 2 (eef-2), commonly used housekeeping genes that all show low Gini coefficients and high bias terms, supporting the hypothesis that bias term analysis can be directly used for housekeeping gene identification.

### Master regulator identification

Co-expressed genes often share similar transcription factor binding sites (TFBSs) in their promoter, and the degree of co-expression is correlated with the number of shared TFBSs^25^. The identification of shared transcriptional regulators of the genes influenced by one neuron should thus allow the identification of master regulators of both gene sets and cell types. To investigate this, we ran our network on a scRNA-Seq dataset of 2,552 cells from mouse testis^26^. We obtained gene sets from our model by extracting the mappings from decoder hidden layer neurons to genes and performed sequence motif enrichment analyses using HOMER^27^. From these, we identified gene sets targeted by the transcription factors CREMt, MYBL1, and MEF2A. As previously described, we found the CREMt motif to be strongly associated with gene sets that were activated by cell type specific latent variable space neurons corresponding to late stages of spermatogenesis, but not to stages before the completion of meiosis^28^ (Fig. 3a). In contrast, the motif that MYBL1 is known to bind to was strongly enriched in gene sets that were specifically activated in pre- and early meiotic cells, peaking around prophase I (Fig. 3b), also in line with previous literature describing MYBL1 to be a master regulator of meiosis^29^. A single neuron was identified that showed the MEF2A associated motif as top enriched TFBS. MEF2A has been described to be restricted to somatic cell populations in the testis^30^. Inspection of the weight mappings confirmed the hypothesis that the associated gene set should thus be most strongly active in Sertoli and Leydig cells (Fig. 3c).

**Figure 3.**
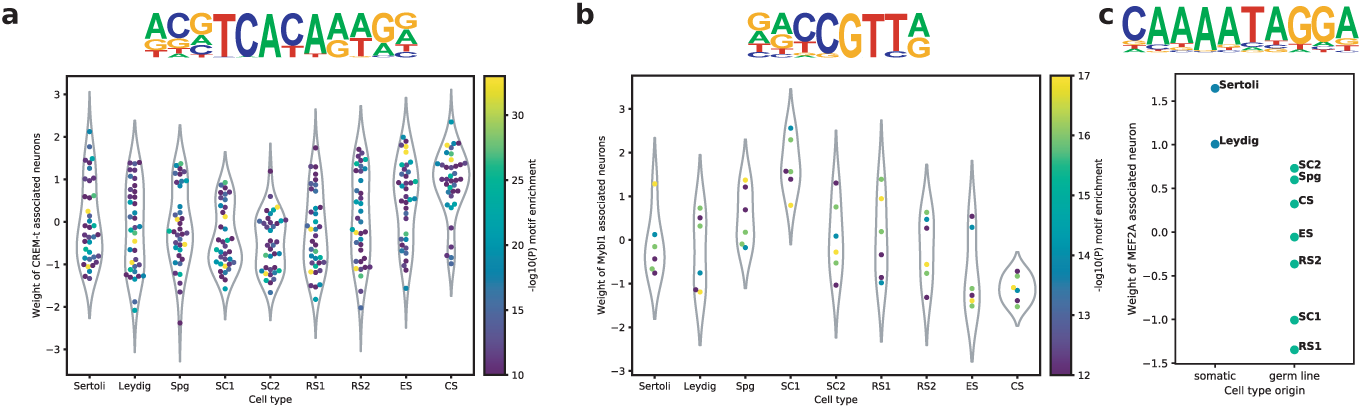
Upstream regulator identification in mouse testis data. a, b) Weight mapping strength from the cell type specific latent variable space (x-axis) to gene set neurons in the first decoder layer that show specific enrichment of a) CREM-t or b) Mybl1 target genes. The motif enrichment of each gene set is color-coded (yellow = lower P-value). The corresponding motif is indicated above the plot. c) Weight mappings as in a) and b) of the single neuron showing a strong enrichment in MEF2A target genes. The x-axis position indicates whether a cell type is somatic. Spg = spermatogonia, SC = spermatocytes, RS = round spermatids, ES = elongating spermatids, CS = condensing and condensed spermatids.

### Batch effect correction

Reserving dimensions of the latent variable space for specific cell types and the introduction of a bias term on the output layer allowed us to isolate specific gene expression programs for individual cell types (Fig. 2a-c). We thus hypothesized that it should be possible to isolate the effects of further characteristics, such as experimental batch, sex, or age. To demonstrate this, we obtained scRNA-Seq datasets of pancreatic islet cells, generated using four differenttechnologies (CelSeq: GSE81076, CelSeq2: GSE85241, Fluidigm C1: GSE86469, SMART-Seq2: E-MTAB-5061)^31–34^. Cell type assignments were obtained from the accompanying datasets of Stuart *et al*.^8^. We restricted the analysis to the four most common cell types (alpha, beta, acinar, and ductal cells) to avoid issues with low cellular coverage of rare cell types in some datasets. The identity matrix for latent variable space assignment was created by concatenating two one-hot encoded matrices, one for cell type and one for technology. For comparison purposes, a second model was trained with the cell type as only assignment variable for the latent variable space, leaving all other parameters and hyperparameters identical. We then evaluated the quality of batch correction by comparing the mappings from the cell type specific latent variable space to specific markers genes for each cell type. We chose *CPB1* as marker for acinar cells, glucagon (*GCG*) for alpha cells, insulin (*INS*) for beta cells, and *CFTR* for ductal cells. The weights observed in the batch corrected model were far more strongly associated with the correct cell type when compared to the model without batch correction (Fig. 4a). The list of genes with strong mappings from the batch or technology part of the latent variable space was dominated by ERCC spike-in controls and mitochondrial transcripts: of the 20 genes with the highest weight mappings to the batch dimensions, six were mitochondrial genes, and four were ERCC spike-ins. The percentage of mitochondrial genes is frequently used as an indicator of the quality of cells, as mitochondrially transcribed RNA is retained even when cytoplasm is lost in broken cells^35^. Finding the expression of these genes attributed to batch effects is thus unsurprising, as different single-cell isolations are likely to be of differing quality, even more so when they are performed with complex protocols and at different labs. Of note, known markers for individual cell types in the pancreas had a strong mapping to the batch dimension. Investigation of these markers, *PCSK1N* (ranked 2^nd^ in batch weight) and *GCG* (ranked 6^th^ in batch weight) revealed that they had been correctly identified as genes associated with batch effect. Despite being described as excellent marker for endocrine cells, *PCSK1N* expression was almost exclusively found in the Smart-seq2 dataset^34,36^ (Fig. 4b). As the authors of the corresponding study found a correlation of *PCSK1N* and BMI, this observation may be due to differences in the patient collective. The high weight mappings of *GCG* were caused by the presence of this transcript in all cell types in the Smart-seq2 dataset, while being absent from non-alpha cells in the other datasets (Fig. 4c). This might be caused by contaminating RNA molecules from broken alpha cells in this dataset.

**Figure 4.**
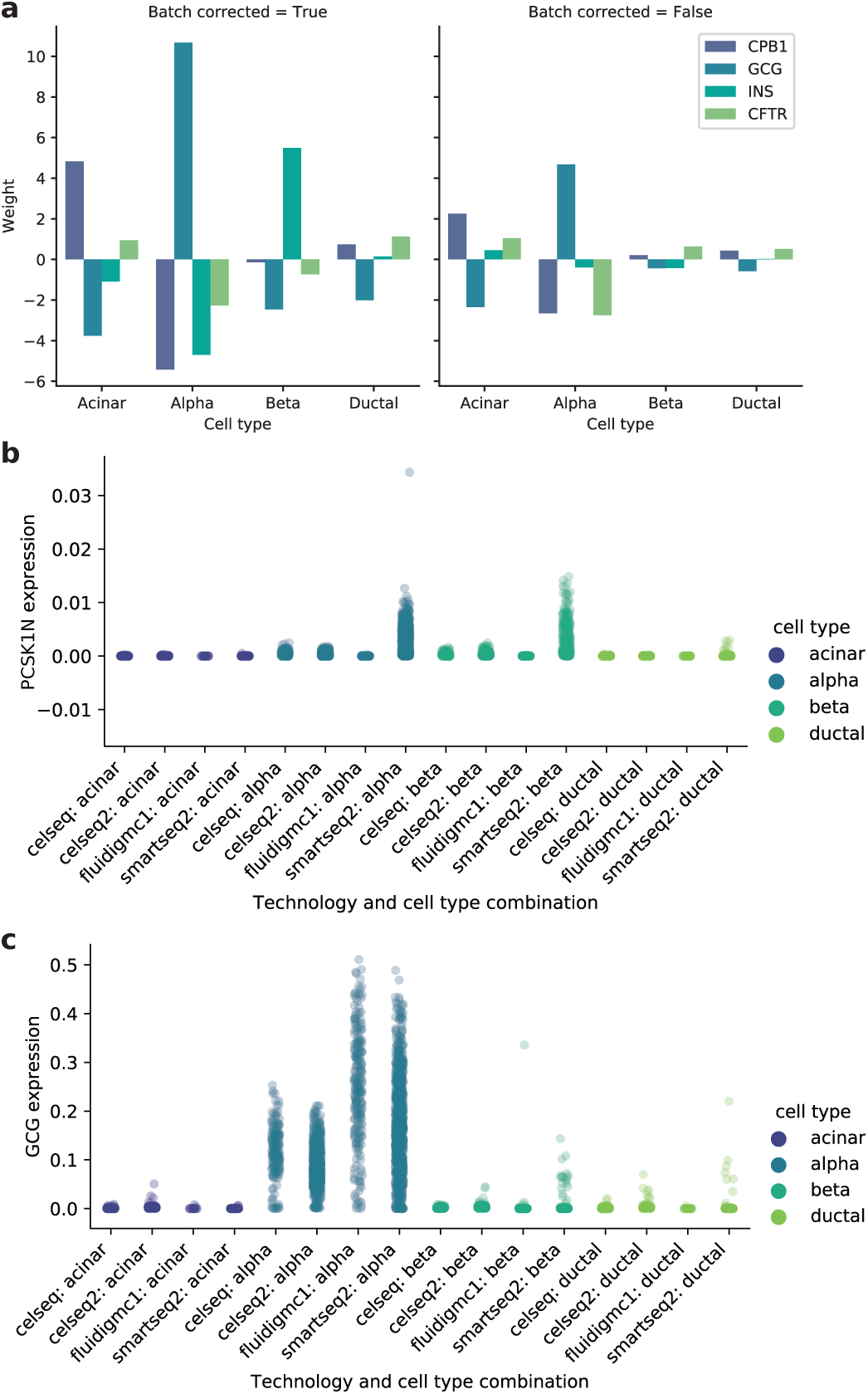
Batch effect correction in human pancreatic islet data from different sources. a) Weight mappings from the cell type specific latent variable space to four marker genes. Left panel: batch correcting network encoding the batch in the latent variable space. Right panel: standard network without batch correction. b, c) Normalized expression values of b) glucagon and c) PCSK1N in different cell types and techniques.

## Discussion

It should be noted that gene sets identified by our model are not ordered and may be redundant to some extent. While this redundancy can be reduced by imposing stricter penalties on the L1 norm, manual inspection of the gene sets reported is still necessary. The same holds true for matrix factorization based techniques, which have however been successfully used for the analysis of and hypothesis generation based on scRNA-Seq^10^. Compared to these methods, our algorithm has some major advantages. Firstly, it allows for the direct identification of gene sets active in a specific cell type without having to infer this through population averages. Secondly, batch effects can be isolated from other contributors to gene expression. With standard techniques based on NMF, a gene set’s activity might not be clearly attributable to a batch effect, as no separation between different variables is enforced.

In summary, we present a conditional variational autoencoder layout based on a combination of variational autoencoders and matrix factorization techniques that can be utilized to identify gene modules such as co-expressed gene sets and pathways. Reserving dimensions of the latent variable space for specific parameters of our input datasets allows us to identify gene sets specific for certain cell types and isolate batch effects and other confounders. The identification of static components of gene expression, i.e. housekeeping genes, is possible through the inspection of a bias term in the linear part of the model. The presence of shared transcription factor binding sites in the promoters of identified gene sets provides support for the notion that the identified relationships have a biological relevance rather than being spurious correlations.

Although this study concentrates only on gene expression studies, CVA is not restricted to this particular application. As it is conceptually similar to deep NMF models, it can aid the analysis of any form of data that can be structured, interconnected, and overlapping sets in other fields such as natural language processing or market segmentation.

## Methods

### Data preprocessing

Example datasets were obtained as described in the data availability section. Genes that were expressed in less than three cells were removed from the expression matrices. No further filtering was performed. Cluster assignments were obtained from the respective download sources.

### Input formatting

Our model has two input requirements. The first is a matrix **X** ∈ ℝ^*n*×*m*^ with *n* cells and *m* genes, where *x*_*i,j*_ denotes the expression values of gene *j* in cell *i*. The second input is a vector of cluster identities ***y*** ∈ ℕ < *c*, where c is the number of cell types with *y*_*i*_ denoting the cluster identity of cell *i*. In an initial step, each row of the expression matrix is scaled to unit norm, with ‖*x*_*i*_‖_1_ being the *ℓ*_2_ norm of cell *i*:

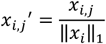

Furthermore, in the case of a single cell type variable, e.g. hard clustering, the vector of clusters ***y*** is one-hot encoded, yielding a binary matrix of shape *n* × *c*. Note that in case of mixed cell identities, such as doublets or soft-clustering results, decimal numbers can be used instead of zeros and ones to indicate mixing proportions. If several variables are to be encoded, the corresponding matrices can be concatenated, leading to a matrix of shape *n* × (*c*_2_ + *c*_9_ + … + *c*_*k*_) for *k* classes.

### VAE encoder

The first two layers of the model are standard densely connected layers of 128 and 64 neurons, respectively, with rectifier linear unit (ReLU) activation functions:

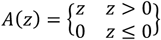

The layer number and size worked well for the example datasets presented here, but the option to alter/specify the values remains in our implementation. The dense layers are followed by two layers representing the mean and log variance of a normal distribution. This allows to model hidden influence factors in a way that emphasizes the mean while retaining variation, rather than reducing the networks loss by outputting the mean^19,20^. Furthermore, a constraint is placed on these two layers to minimize the Kullback-Leibler divergence to a Gaussian *𝒩*(0 + *s*; 1), where *s* is a location shift parameter:

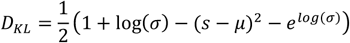

This allows generating simulated data by sampling from a Gaussian *𝒩*(0 + *s*; 1) and passing the values to the decoder part of the network. While in standard VAEs *s* is 0, we use a sufficiently large value (>= 10, equal to 10σ) to shift the mean of the Gaussian away from 0 and avoid mixing with the zero values obtained due to the enforced sparsity for inapplicable classes (see below).

In contrast to regular implementations of variational autoencoders, the following is applied for latent index *l* ∈ {ℤ × *c*} of a cell with cluster identity *k*:

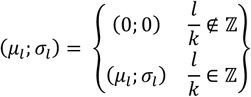

The same is applied to the Kullback-Leibler loss, effectively just considering a reserved subspace of the latent dimension for cell attributed to each cluster. In case of partial cell identities, all cluster subspaces are scaled according to cell type proportions.

The actual latent layer *z* is then calculated from the mean and the log variance as^19^:

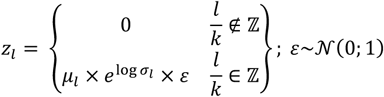

### VAE decoder

In the decoder part of the network, the output of this latent layer is then fed into two densely connected layers of 256 and 512 neurons with ReLU activation, batch normalized, and passed to the final layer with *m* neurons and a softmax activation function:

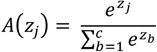

The latter ensures all expression values of a cell sum to one, which is also true for the scaled input.

The cost function of the network is the sum of the Kullback-Leibler divergence and the mean-squared error of the reconstruction:

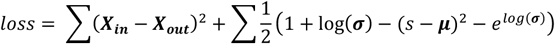

In our implementation, the mean-squared reconstruction error can be changed to mean absolute error, if large-scale errors are not considered to be relatively more important.

Since our model is run to identify gene co-expression modules, the hidden layers of the decoder are used without biases, simplifying their activation from:

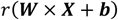

where *r* is the ReLU function, ***W*** is the weight matrix, ***X*** are the inputs, and ***b*** are the biases, to:

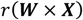

In order to arrive at distinct gene co-expression modules, a dynamic weight regularizer is employed in this mode imposing a penalty of:

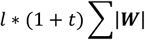

where *l* is a scaling factor for the penalty and *t* is the training epoch. Again, as these can be critical tunable hyperparameters, our implementation allows the user to choose either a dynamic or static *L*_1_, *L*_2_, or combined regularizer with a user-defined initial penalty *l*.

### Obtaining gene sets from weight matrices

The strength of connection between a hidden decoder layer neuron and an output gene is then estimated as the sum over all weights from this neuron through the following layers to the output, disregarding the ReLU function:

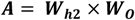

Where 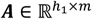 is an activity matrix with *h*_*1*_ as the size of hidden layer 1 and *m* as the output size (gene number), 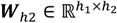 is the weight matrix of hidden layer 2, and 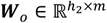 is the weight matrix of the output layer. If more than two layers are involved, the matrix multiplication is carried out step by step to arrive at the final activity matrix.

In order to identify the top associated genes per neuron in the activity matrix, the elbow point for positive and negative enrichment is calculated. The elbow point in this case is the point of maximum distance from the diagonal connecting the lowest and highest values on a sorted list of weights. When scaling the weights and ranks of the positive or negative weights to the range [0,1), this diagonal has a slope of 0 and an intercept of 1, so *y* = *x* + *n*. The distance between the curve of weights and the diagonal is calculated along a line perpendicular to the first diagonal, going through the point on the weight curve (*x*_*w*_, *y*_*w*_) with an intercept *n*_*w*_, so *y*_*w*_ = −*x*_*w*_ + *n*_*w*_ ⟺ *n*_*w*_ = *y*_*w*_ + *x*_*w*_. The intersect of both diagonals is at 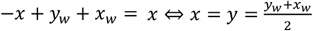. It follows that the distance of the weights curve from the diagonal is 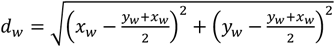. Since *x*_*w*_ and *y*_*w*_ are both strictly positive and *x*_*w*_ ≥ *y*_*w*_, this simplifies to *d*_*w*_ ∝ *x*_*w*_ − *y*_*w*_.

### Housekeeping gene identification

In contrast to the hidden decoder layers, the bias vector is retained as trainable parameter in the calculation of the output layer activation, capturing the static components of gene expression, yielding the following activation:

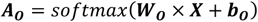

*b*_*o*_ is a vector whose length equals the number of genes, indicating a sort of steady-state activity of that gene irrespective of the variables encoded in the latent variable space.

### Batch effect correction

For batch effect correction, the latent variable space of the network was constructed as described above, assigning latent dimensions to batches as well as to cell types. The weight mappings from the batch specific latent variable space were then analyzed in isolation from the cell type specific ones. In cases were smoothing and batch effect correction are desired, one could train the network as described here and then perform prediction with a class matrix whose batch components are set to zero.

### Gene set enrichment

Gene set enrichment calculations were performed using Metascape^24^. The set of genes annotated in the expression matrix before filtering was used as background.

### Motif enrichment

Motif enrichment was performed using the findMotifs.pl command of HOMER v4.10^27^. The standard promoter annotations for the mm10 mouse genome as of July 2019 were used. Genes contained in the filtered expression matrix, but not enriched in a specific gene set, were used as background.

## Data availability

All example datasets were used in previously published studies. The *C*. *elegans* dataset was downloaded according to the instructions provided at http://atlas.gs.washington.edu/worm-rna/docs/#use-case-1-expression-pattern-of-a-gene-of-interest^23^. The testis scRNA-Seq data with the accession number GSE104556 was downloaded from GEO^26^. The set of pancreas scRNA-Seq datasets including annotations was downloaded according to the instructions on https://satijalab.org/seurat/v3.0/integration.html. The individual datasets can be accessed on GEO (GSE81076, GSE85241, GSE86469) and SRA (E-MTAB-5061)^31–34^.

## Code availability

The code for our implementation of the CVA algorithm is available at https://bitbucket.org/conrad_lab/CVA.

## Acknowledgements

This publication is part of the Human Cell Atlas - www.humancellatlas.org/publications.

## Author Contributions

SL and CC conceived this study. SL implemented the algorithm and analyzed the data. FWT assisted with the implementation. CC and RE supervised the study.

## Competing interests

The authors declare no competing interests.

## Notes

#### Summary of Updates

Acknowledgements updated

